## Supplementary figures for "Extrachromosomal DNA is associated with chromothripsis events and diverse prognoses in gastric cardia adenocarcinoma"

### Supplementary information:

**Supplementary Table 1**: Sequencing quality assessment of whole genomic sequencing (WGS) data from 36 pairs of GCA tumour and adjacent normal tissues.

**Supplementary Table 2**: Detailed characterization of individual ecDNA amplicons from the prediction of WGS data using AmpliconArchitect (AA).

**Supplementary Table 3**: Characterization of oncogene co-amplification in ecDNA amplicons.

**Supplementary Table 4**: Clinicopathological records, microsatellite instability (MSI) staining, and chromosome instability (CIN) grade in 36 GCA tumours.

**Supplementary Table 5**: Sequencing quality assessment of whole exome sequencing (WES)data from 75 pairs of GCA tumour and adjacent normal tissues.

**Supplementary Table 6**: Clinicopathological records in 75 GCA patients with whole exome sequencing (WES)data.

**Supplementary Table 7**: *ERBB2* RNA expression and ERBB2 protein expression in 44 GCA patients.

**Supplementary Table 8**: Clinicopathological records and ERBB2 protein immunohistochemistry (IHC) staining in 1668 GCA patients.

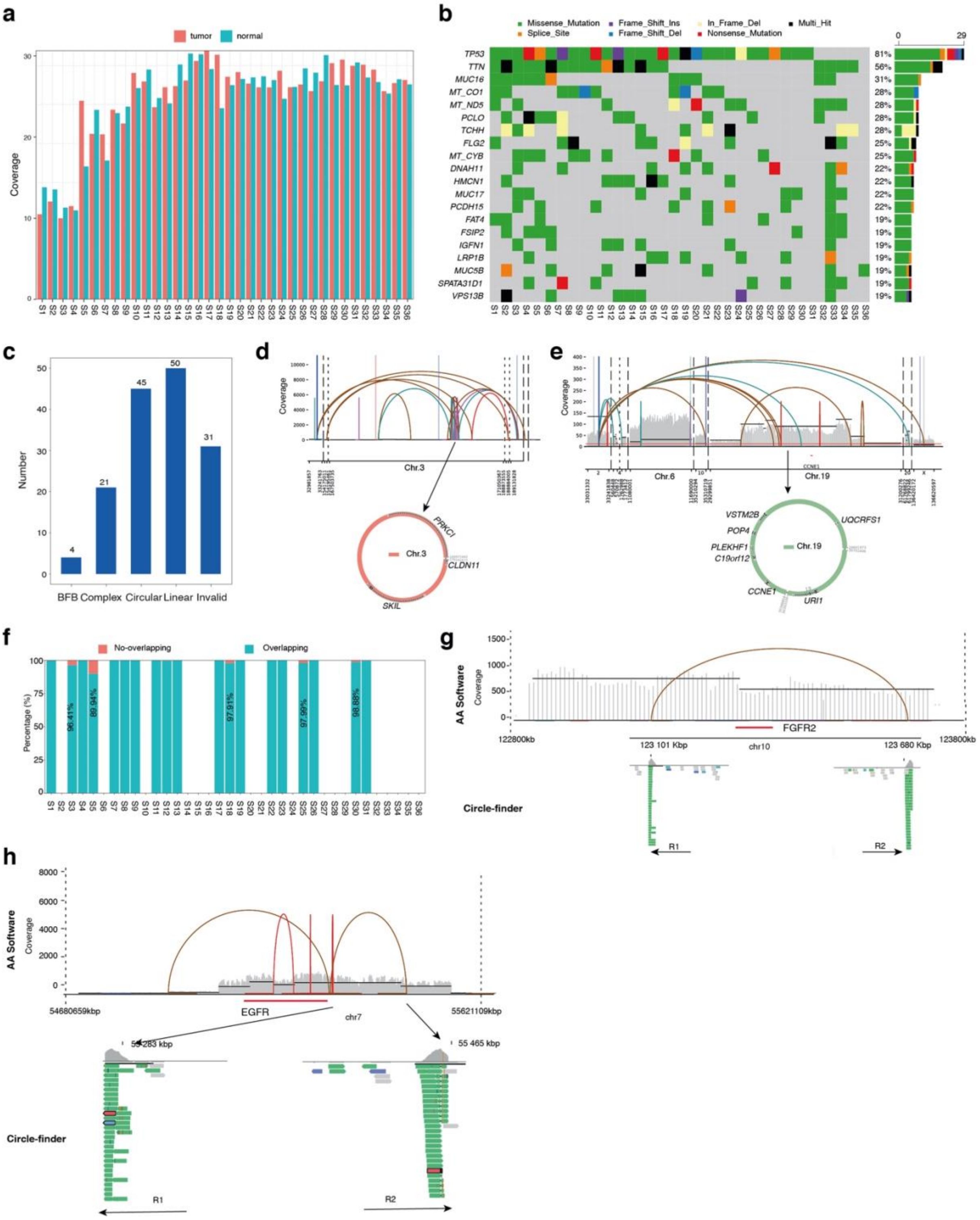


**Supplementary Figure 1**: **Prediction of ecDNA amplicons from the Chinese GCA cohort using whole genomic sequencing (WGS).**

**a**, Sequencing depth of WGS from 36 (S1-S36) pairs of GCA tumour and adjacent normal tissue. Tumour = GCA tumour tissue; Normal = adjacent normal tissue.

**b**, Mutation frequency of genes from 36 (S1-S36) GCA patients.

**c**, Number of different types of ecDNA amplicons predicted by AmpliconArchitect (AA) software in the 36 GCA cohort, where ecDNA amplicons were further classified into circular, complex, linear, breakage-fusion bridge (BFB) and invalid.

**d and e,** Examples of circular ecDNA amplicons predicted by AmpliconArchitect (AA) software and constructed into circular format.

**f**, Summary of overlapping frequency in circular ecDNA amplicons from prediction of AmpliconArchitect (AA) and detection using Circle-finder. The empty bar represents no ecDNA or no circular ecDNA amplicons identified by AA.

**g, h**, Example of circular ecDNA amplicons detected by both AA software and Circle-finder. Bottom panel: The location of forward and reverse sequencing reads (R1 = read 1, R2 = read 2) in the circular junction point are indicated on the genome browser.

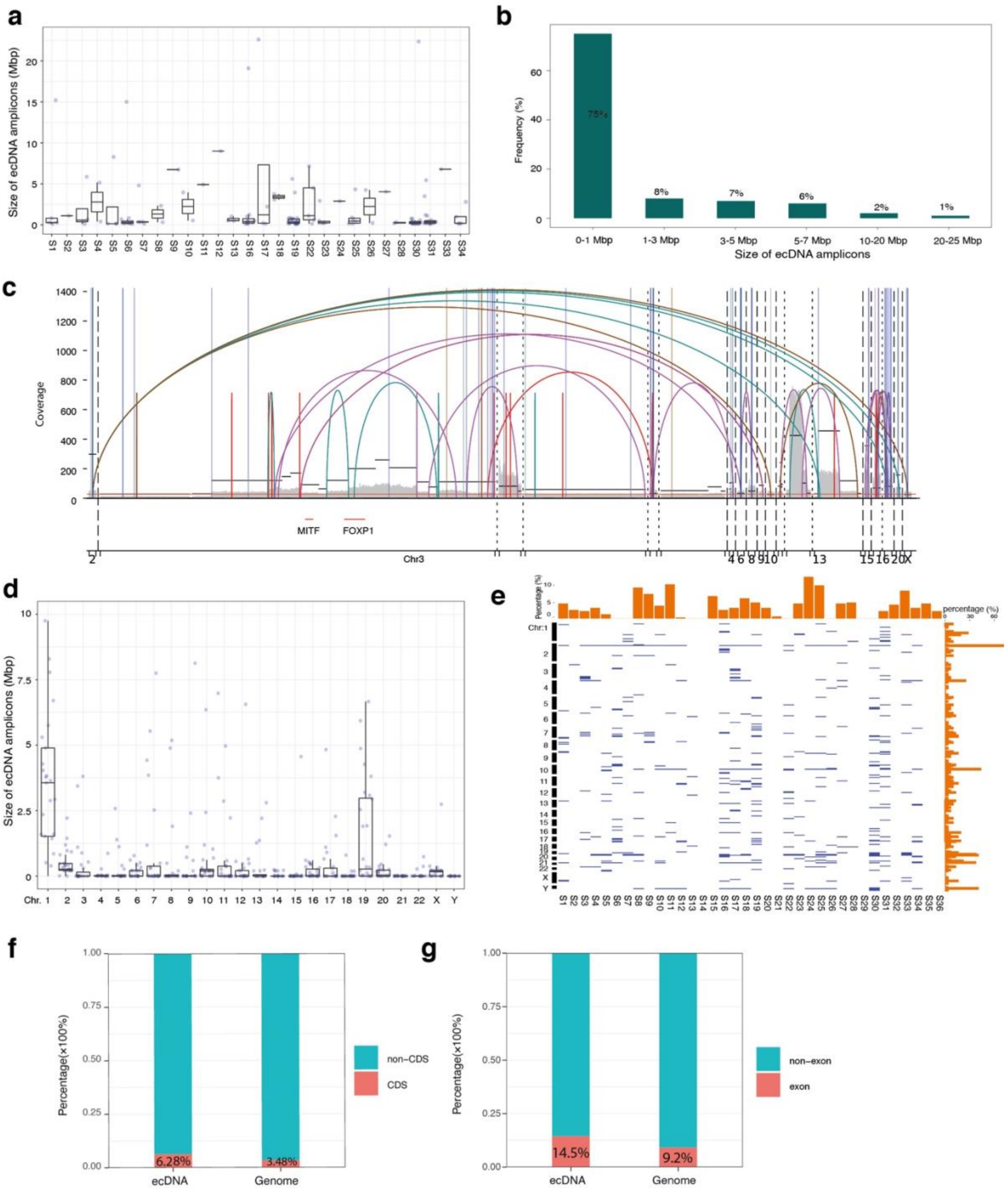


**Supplementary Figure 2: Characterization of ecDNA amplicons in the GCA cohort.**

**a,** Summary of ecDNA amplicon size in each GCA patient, where each dot represents one ecDNA amplicon.

**b,** Size distribution of ecDNA amplicons in our GCA cohort.

**c**, Example of large ecDNA amplicons (> 20 Mbp) deconvoluted into multiple potential combinations of amplicons using AA software, where different connection lines on the top represent potential combinations within ecDNA amplicon regions.

**d,** Summary of ecDNA amplicon size in each chromosome from the cohort. Each dot represents one ecDNA amplicon.

**e,** Distribution of ecDNA amplicons in each chromosome and each patient in the cohort.

**f**, Occupancy comparison of coding sequence (CDS) and noncoding sequence (non-CDS) regions of ecDNA amplicons and the whole genome, where the percentage was calculated as follows: length of CDS or non-CDS regions divided by the total length of ecDNA amplicon regions or whole genome.

**g,** Occupancy comparison of exon and non-exon regions of ecDNA amplicons and the whole genome, where the calculation strategy is the same as in **f**.

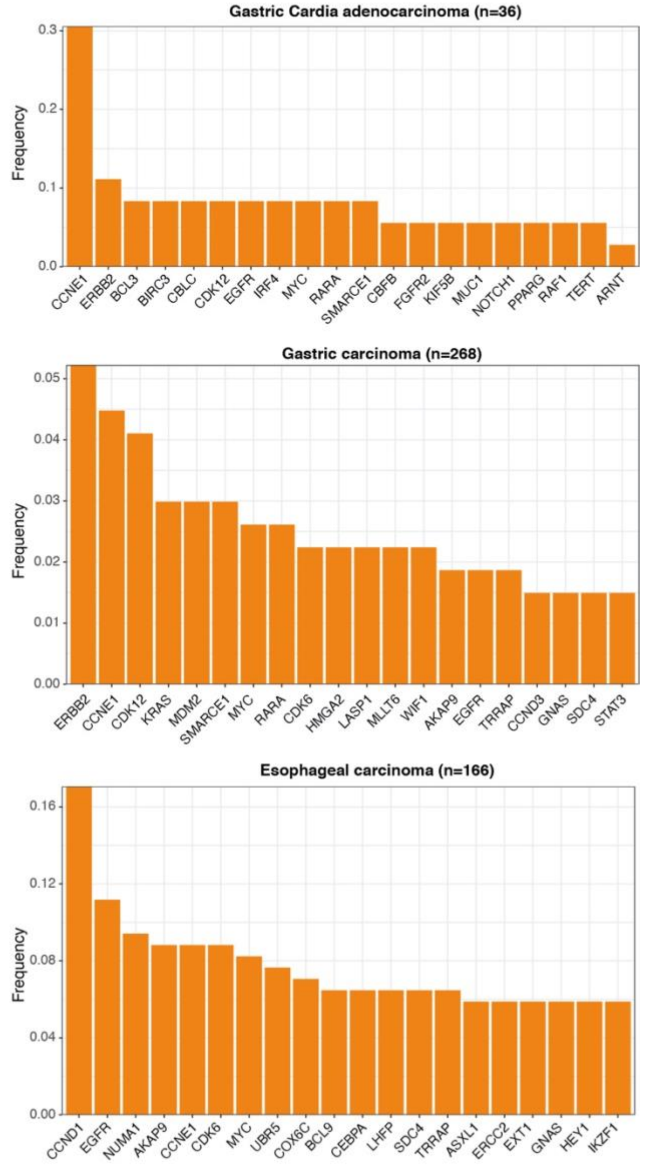


**Supplementary Figure 3**: **Comparison of lists of oncogene or tumour suppressor gene ecDNA amplicons detected in our GCA cohort, TCGA gastric carcinoma and TCGA esophageal carcinoma.**

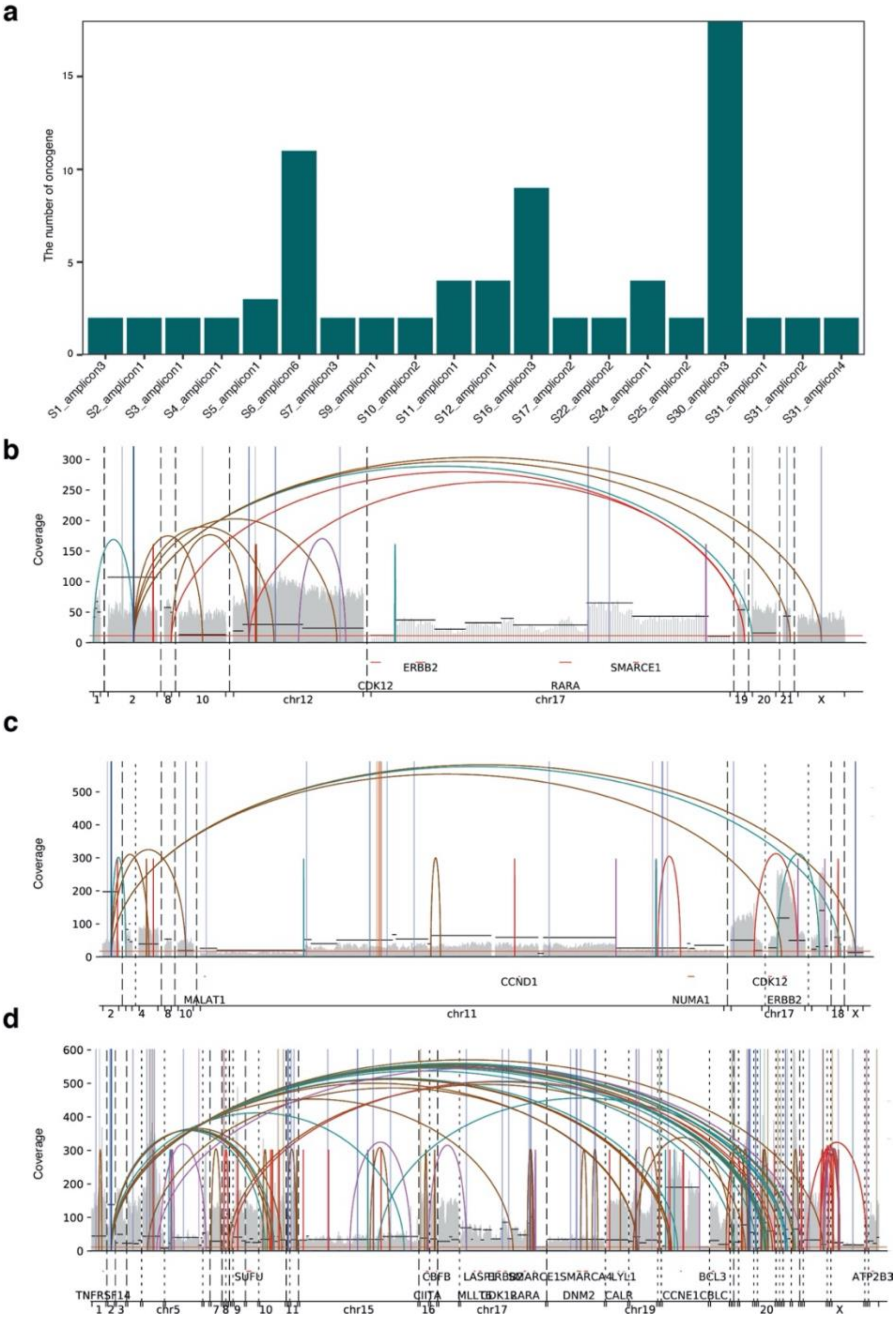


**Supplementary Figure 4**: **Oncogene ecDNA coamplification in GCA.**

**a,** Number of oncogenes and tumour suppressor genes in each ecDNA amplicon. S = Sample. Examples of oncogene and tumour suppressor gene ecDNA co-amplification in our cohort: **b** (including the following genes: CDK12, *ERBB2*, *RARA*, and *SMARCE1*)**, c** (including *the* following genes: *CCND1*, *NUMA1*, *CDK12* and *ERBB2*)**,** and **d** (including the following genes: *ERBB2*, *RARA*, *CCNE1*, *LYL1*, *CDK12*, *CIITA*, *MLLT6*, *DNM2, TNFRSF14, CBFB, SMARCE1, SUFU, LASP1, CBLC, ATP2B3, SMARCA4, CALR, BCL3*).

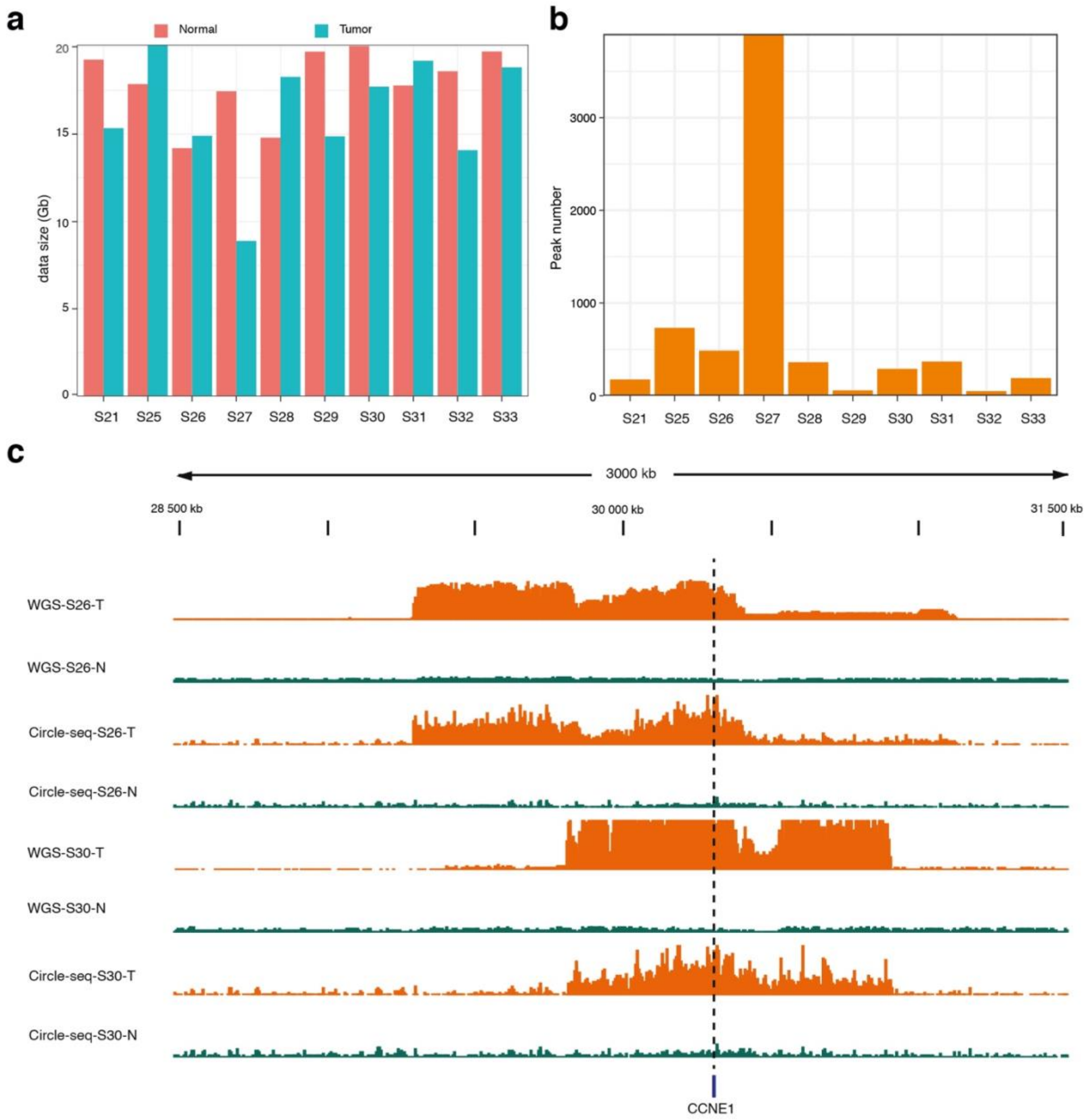


**Supplementary Figure 5**: **Validation of ecDNA amplicons using Circle-seq**.

**a**, Sequencing coverage summary of Circle-seq from GCA patients (n = 10). Tumour = GCA tumour tissue; Normal = adjacent normal tissue.

**b**, Summary of peak numbers from Circle-seq for each GCA patient. Each peak represents one DNA fragment in the circular DNA.

**c**, Genome browser track at the *CCNE1* gene from whole genome sequencing (WGS) and Circle-seq. The dotted line indicates the location of the *CCNE1* gene. N = adjacent normal tissue, T = GCA tumour tissue.

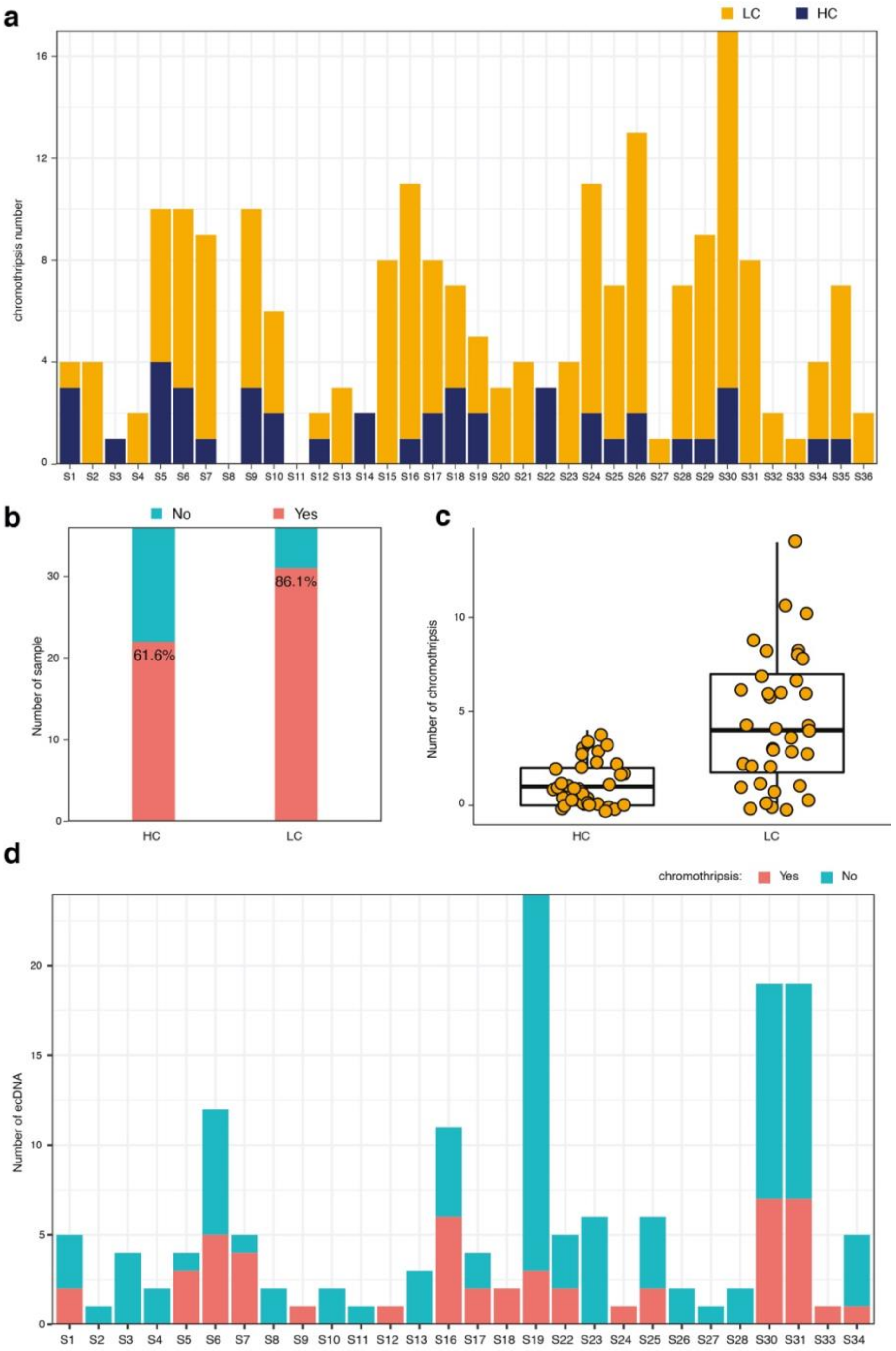


**Supplementary Figure 6**: **Detection of chromothripsis in the GCA cohort.**

**a,** Summary of predicted chromothripsis events in the GCA cohort using different parameters. HC = high confidence chromothripsis, LC = low confidence chromothripsis.

**b,** Frequency of chromothripsis detected in the GCA cohort. HC = high confidence chromothripsis, LC = low confidence chromothripsis.

**c**, Range of chromothripsis events detected from the high confidence (HC) and low confidence (LC) samples in the GCA cohort.

**d**, Summary of chromothripsis events occurring at regions of ecDNA amplicons in our GCA cohort.

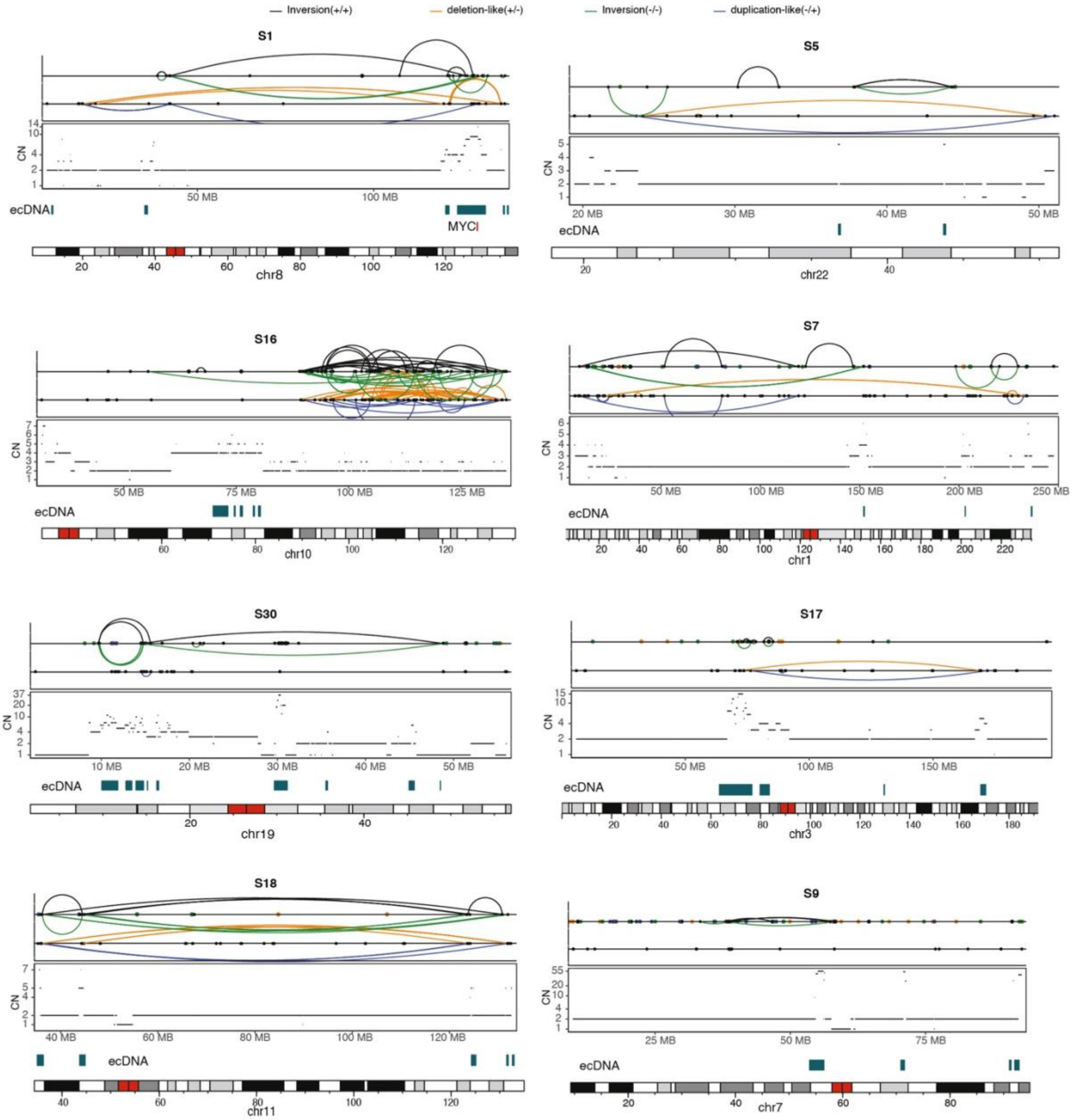


**Supplementary Figure 7**: **The fine structure of detected chromothripsis at different ecDNA amplicon regions in different GCA patients.** S = Sample**.**

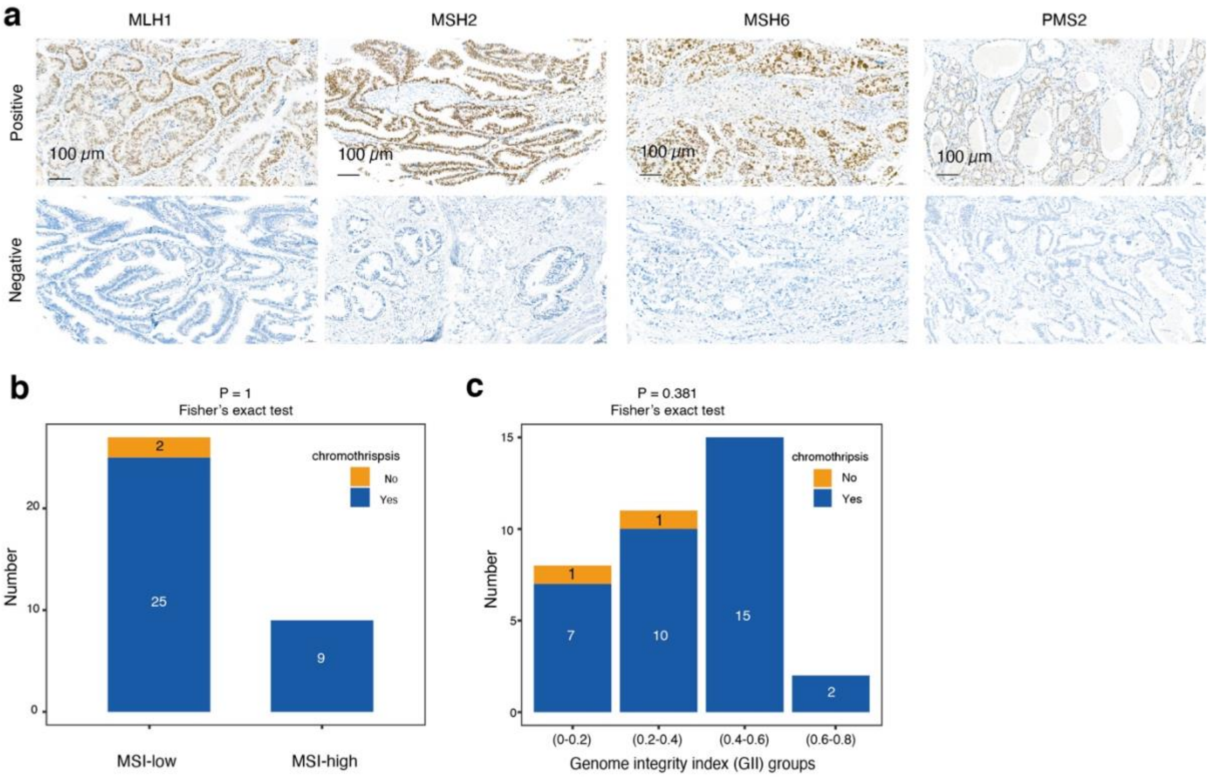


**Supplementary Figure 8**: **Microsatellite instability (MSI), chromosome instability (CIN) and chromothripsis events.**

**a**, Representative immunohistochemistry (IHC) images of four proteins (MLH1, MSH2, MSH6 and PMS2) from 36 patients. If one of four proteins exhibited negative staining by IHC, we labelled the patient as MSI-high. If all four proteins exhibited positive staining by IHC, we labelled the patient as MSI-low.

**b**, Presence and absence of chromothripsis events in the MSI-high and MSI-low groups of GCA patients. The numbers on the bars are patient numbers.

**c**, Comparison of chromothripsis events in different groups of chromosome instability (CIN), where CIN was divided into 4 groups based on different genome integrity indices (GIIs). The numbers on the bars are patient numbers.

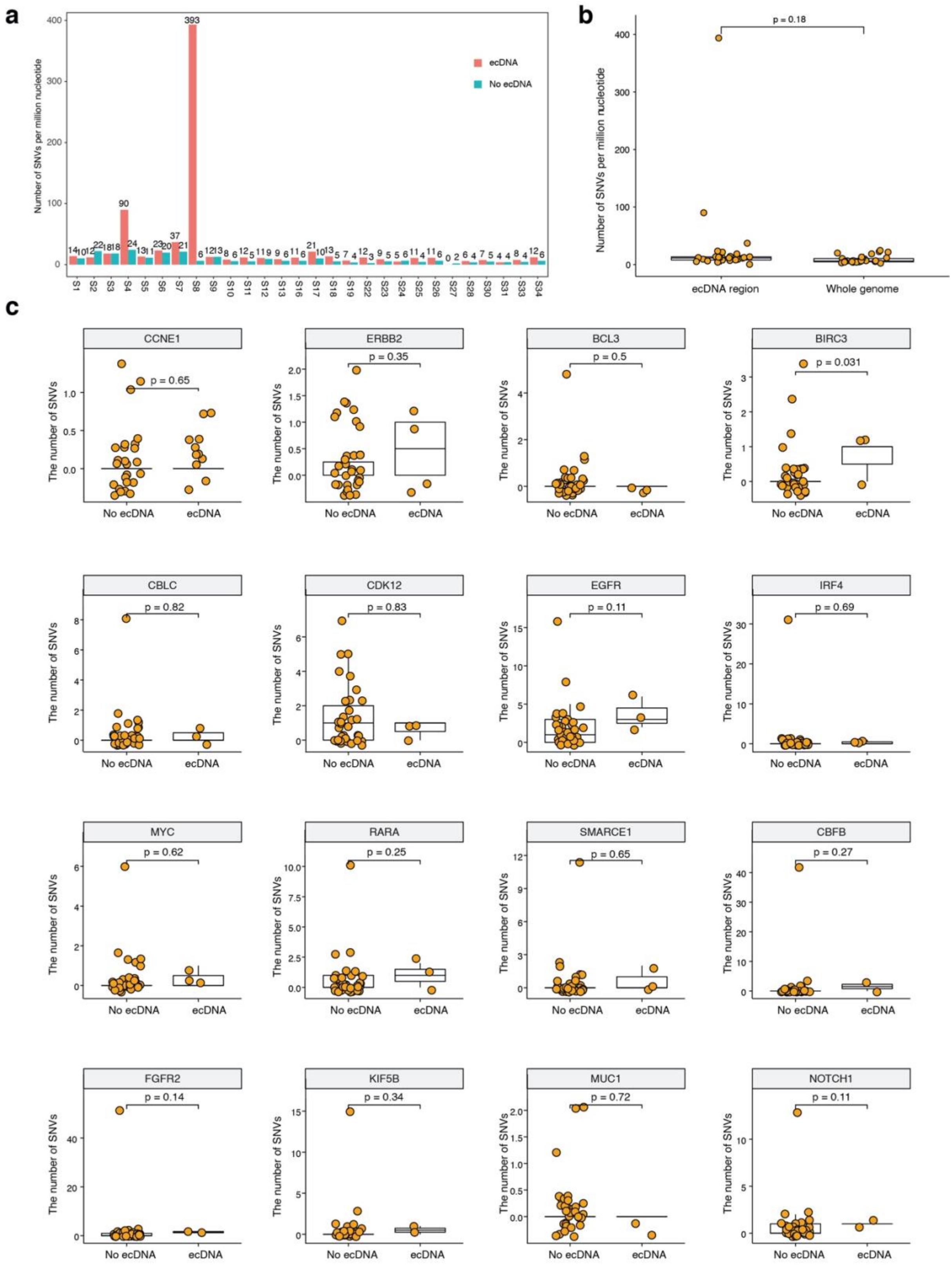


**Supplementary Figure 9**: **Somatic gene mutation and ecDNA amplicons**.

**a**, Comparison of the average number of single-nucleotide variants (SNVs) per million nucleotides from regions of ecDNA amplicons and from the whole genome in individual patients. S = Sample.

**b**, Comparison of the average number of single-nucleotide variants (SNVs) per million nucleotides from regions of ecDNA amplicons and from the whole genome in all 36 patients. Each dot represents one patient. The p-value was calculated using the Wilcoxon signed-rank test.

**c**, Comparison of SNV numbers on oncogenes or tumour suppressor genes between the ecDNA amplicon-present and ecDNA amplicon-absent groups in the cohort, where 16 genes (oncogenes or tumour suppressor genes) were selected, and ecDNA amplicons were observed in at least two patients. Each dot represents one sample. The p-value was calculated using the Wilcoxon signed-rank test.

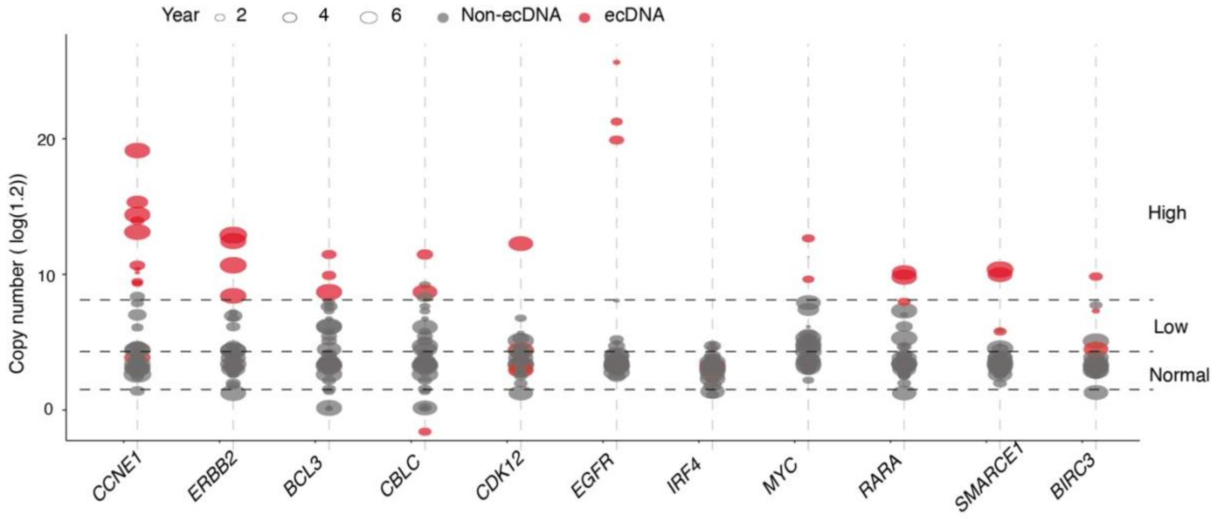


**Supplementary Fig. 10**: **Relationship between oncogene or tumour suppressor gene amplification, the presence of oncogene ecDNA amplicons and patient prognosis in 36 GCA patients.** The copy numbers of oncogenes or tumour suppressor genes were divided into three groups: High, Low and Normal. High = high copy number of gene amplification, Low = low copy number of gene amplification, Normal = no gene amplification.

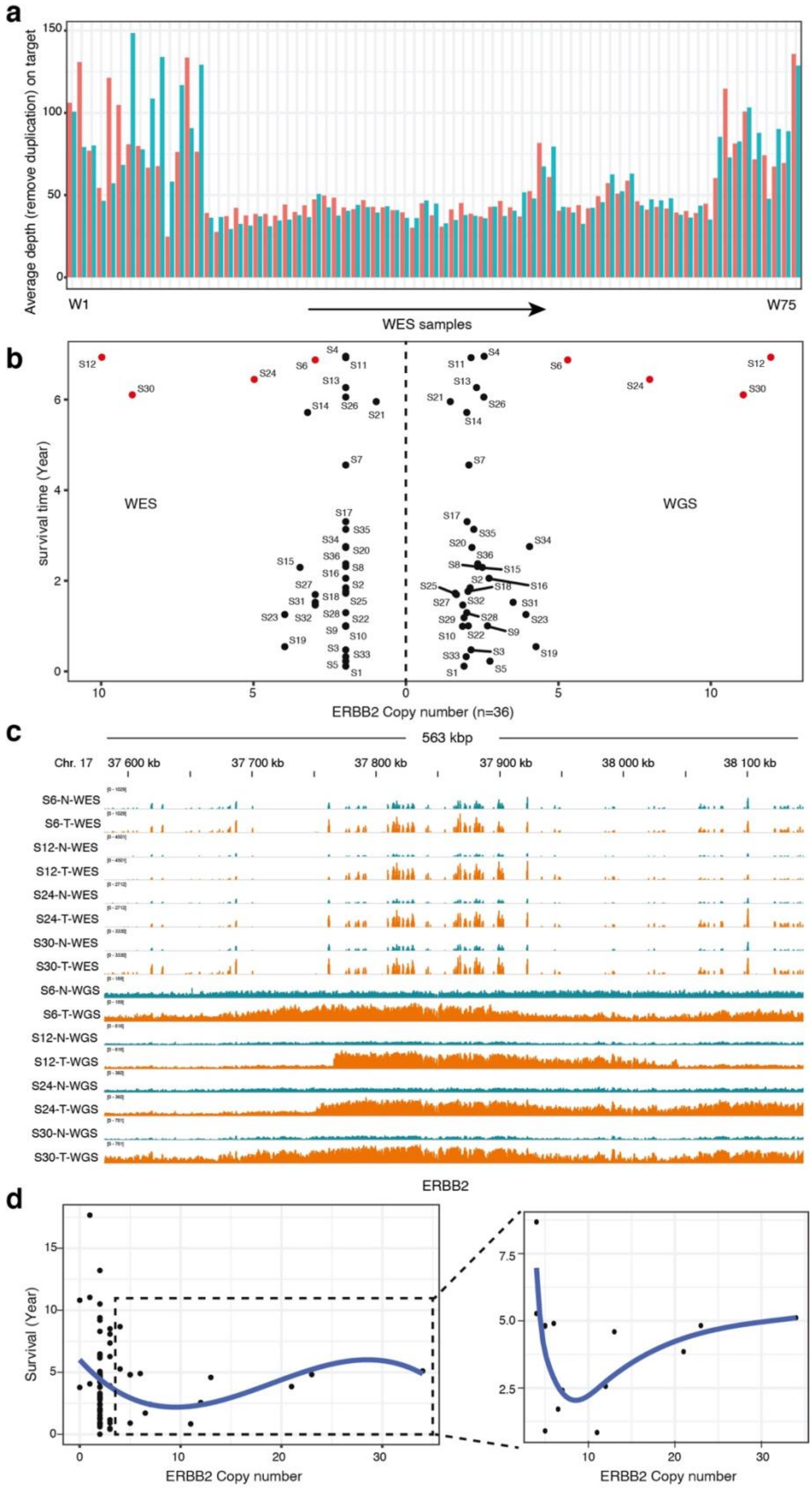


**Supplementary Figure 11**: **Correlation study of gene copy number from whole exome sequencing (WES) and patient prognosis**.

**a**, Sequencing coverage of WES from 75 pairs of GCA tumour and matched adjacent normal tissue. Tumour = GCA tumour tissue; Normal = adjacent normal tissue. S = Sample.
**b**, Comparison of the gene copy number of the *ERBB2* gene between WES and whole genome sequencing (WGS) in the same GCA samples. S = Sample.
**c**, Genome browser track of the *ERBB2* gene locus of both WES and WGS data from 4 pairs of GCA patients;-T- = GCA tumour tissue; N- = adjacent normal tissue.
**d**, Correlation of *ERBB2* gene copy number from WES and patient survival time in 75 GCA patients. Each dot represents one patient.

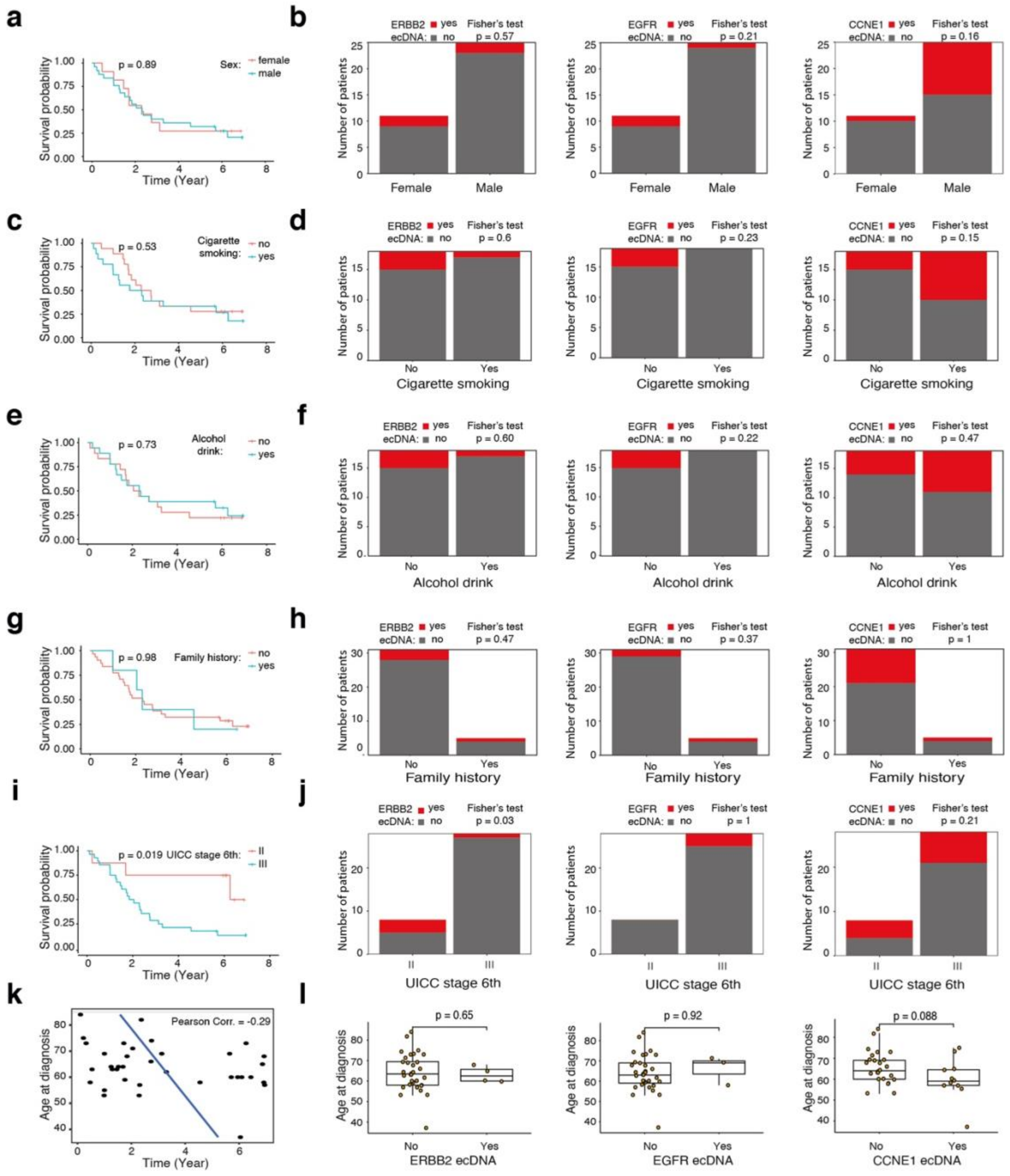


**Supplementary Figure 12**: **Prognostic analysis with clinicopathological variables and the presence of oncogene ecDNA amplicons.**

**a**, Survival analysis of 36 GCA patients based on sex; p-value was calculated using the Log rank test.

**b**, Comparison of oncogene ecDNA (*ERBB2*, *EGFR*, *CCNE1*) presence in different sex groups of 36 GCA patients; p-value was calculated using Fisher’s test.

**c**, Survival analysis of 36 GCA patients based on smoking; p-value was calculated using the Log rank test.

**d**, Comparison of oncogene ecDNA (*ERBB2*, *EGFR*, *CCNE1*) presence in smoking or non-smoking group of 36 GCA patients; p-value was calculated using Fisher’s test.

**e**, Survival analysis of 36 GCA patients based on alcohol consumption; p-value was calculated using the Log rank test.
**f**, Comparison of oncogene ecDNA (*ERBB2*, *EGFR*, *CCNE1*) presence in drinking or non-drinking group of 36 GCA patients; p-value was calculated using Fisher's test.
**g**, Survival analysis of 36 GCA patients based on family history; p-value was calculated using the log rank test.
**h**, Comparison of oncogene ecDNA (*ERBB2*, *EGFR*, *CCNE1*) presence with family history or without family history in 36 GCA patients; p-value was calculated using Fisher's test.
**i**, Survival analysis of 36 GCA patients based on the Union for International Cancer Control (UICC) tumour stage. The p-value was calculated using the log-rank test.
**j**, Comparison of oncogene ecDNA (*ERBB2*, *EGFR*, *CCNE1*) presence in UICC tumour stages II and III of 36 GCA patients. The p-value was calculated using Fisher's test.
**k**, The correlation of survival time and age at diagnosis in 36 GCA patients. Each dot represents one patient.
**l**, The comparison of age at diagnosis of GCA in absent and present oncogene ecDNA (*ERBB2*, *EGFR*, *CCNE1*) groups of 36 GCA patients. The p-value was calculated using the Wilcoxon signed-rank test.

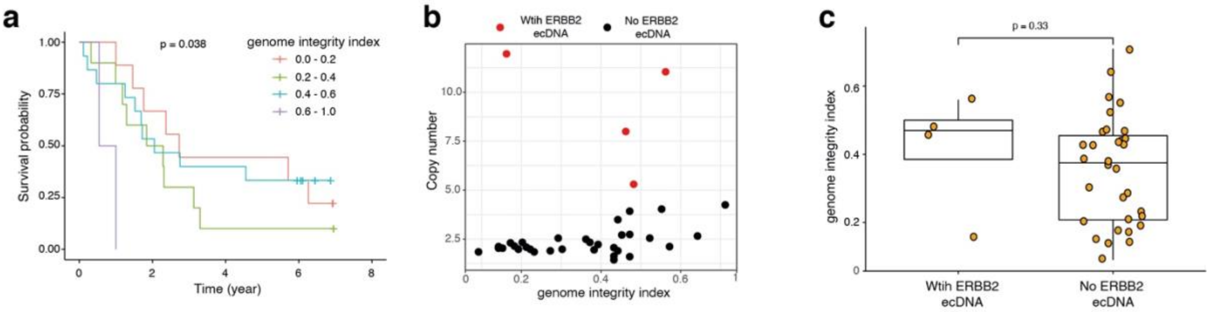


**Supplementary Figure 13: Prognostic analysis based on chromosomal instability (CIN).**

**a**, Survival analysis of 4 groups of CIN defined by the number of genome integrity indices (GIIs). The p-value was calculated using the log-rank test.

**b**, Scatter plot showing the relationship of *ERBB2* gene copy number, presence of *ERBB2* ecDNA (colour-coded) and CIN grade of patients, where each dot represents one patient.

**c**, Comparison of genome integrity index values between *ERBB2* ecDNA present and absent groups. Each dot represents one sample. The p-value was calculated using the Wilcoxon signed-rank test.

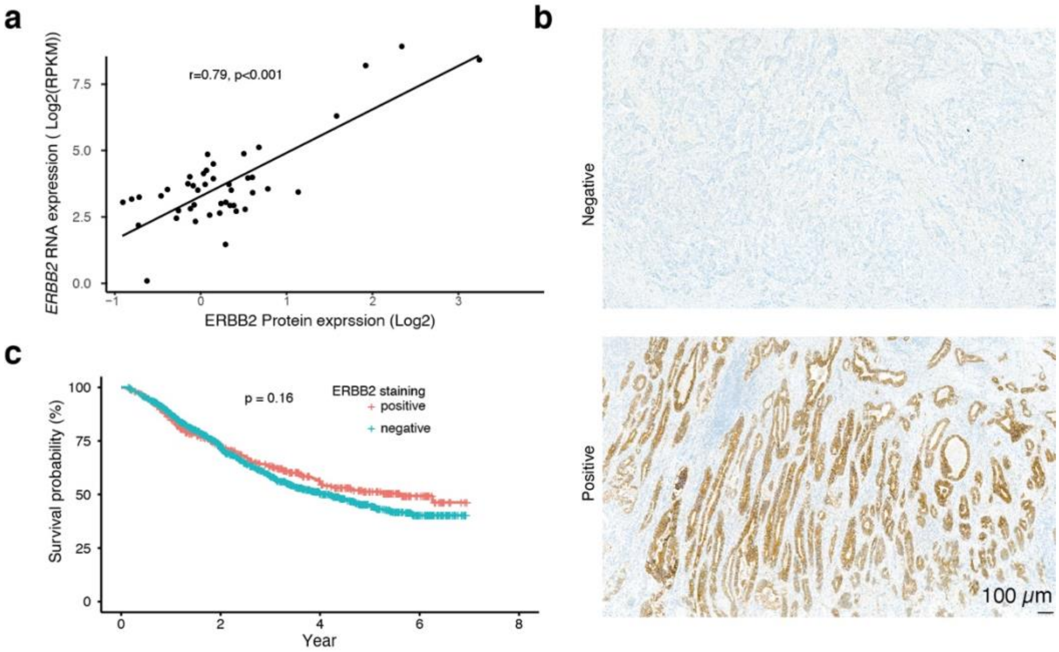


**Supplementary Figure 14**: **ERBB2 protein staining and patient prognosis.**

**a,** Scatter plot showing that there is a positive correlation between *ERBB2* RNA expression level and ERBB2 protein expression level in 44 GCA patients. Each dot represents one patient.

**b,** Representative ERBB2 protein immunohistochemistry (IHC) images of positive (bottom) and negative (top) staining from 1668 GCA patients.

**c,** Survival analysis of positive and negative ERBB2 IHC staining groups in 1668 GCA patients. The p-value was calculated using the log rank test.
